# The eumetazoan origin of PRDM9 function revealed by cnidarian genome analysis

**DOI:** 10.64898/2026.09.07.749791

**Authors:** Amélie Rudler, Laurent Duret, Christoph Grunau, Eve Toulza, Julie A.J. Clément

## Abstract

Where recombination takes place is crucial as it determines the position of genetic reshuffling, which facilitates species evolution and adaptation. In many vertebrates, PRDM9 determines the location of double-strand breaks that initiates meiotic recombination. However, this only applies for *Prdm9* orthologs that possess certain molecular features, such as the presence of the four functional domains (KRAB, SSXRD, PR/SET and ZnF), the conservation of catalytic tyrosines and a fast-evolving zinc finger array. Although the evolutionary origin of PRDM9 has been inferred to the last common ancestor of metazoans, it has been poorly explored outside vertebrates. Here, we explored the genomes of 28 cnidarian species and identified, for all of them, at least one full-length ortholog that appeared to be functional. Striklingly, ten out of the 28 species carried several full-length paralogs with the same molecular markers of functionality. This pattern has never been observed in any other explored taxa. We also identified many truncated paralogs that arose at different times, thereby revealing an important birth-and-death dynamics of *Prdm9* in cnidarians. Phylogenetic analysis revealed that the multiple full-length paralogs are all the result of recent duplication events, occuring mainly after speciation, although truncated paralogs arose all along cnidarian evolution. One of the oldest truncated paralogs appeared before the divergence between Actinaria and Scleractinia. This indicates the probable acquisition of a new function that has been conserved for at least 540 million years and whose nature is still unknown. In addition to providing insights into the ancient origins of PRDM9’s function, our work now raises new questions about the functional redundancy of these multiple full-length paralogs, as well as their evolutionary significance for cnidarian genomes.

## Introduction

Meiotic recombination is a fundamental and evolutionarily conserved process in sexually reproducing organisms across the tree of life. It is initiated by the formation of hundreds of programmed DNA double-strand breaks (DSBs), which are repaired through homologous recombination and lead to the formation of either crossovers (COs), involving reciprocal exchange of genetic material between homologous chromosomes, or non-crossovers (NCOs), which occur without reciprocal exchange (Lam and Keeney 2015). At least one CO per homologous chromosome pair is required to ensure proper chromosome segregation during meiosis (Bolcun-Filas and Schimenti 2012; Borde and de Massy 2013; de Massy 2013).

Meiotic recombination plays also a key evolutionary role by generating novel allele combinations along chromosomes, thereby facilitating adaptation in natural populations (Ritz et al. 2017). In most species, recombination events are concentrated in small chromosomal regions of 1 to 2 kilobases, called recombination hotspots, where the recombination rates can be tens to thousands times higher than in surrounding regions (de Massy 2013). Two mechanisms governing hotspot localization have been described so far. In dogs, birds, plants, and yeast, hotspots preferentially occur in regulatory regions such as promoters and CpG islands, likely reflecting chromatin accessibility and H3K4me3 enrichment (Lichten and Goldman 1995; Muñoz-Fuentes et al. 2011; Pan et al. 2011; Brick et al. 2012; Auton et al. 2013; Choi et al. 2013; Drouaud et al. 2013; Hellsten et al. 2013; Lam and Keeney 2015; Singhal et al. 2015; Stapley et al. 2017). These hotspots tend to be conserved over long evolutionary timescales (Lam and Keeney 2015; Singhal et al. 2015; Kawakami et al. 2017). In contrast, in many vertebrate species, including mice, primates, salmonids, and cattle, recombination hotspots tend to occur outside regulatory regions, and their positions evolve rapidly between closely related species and even between populations (Ptak et al. 2004; Myers et al. 2005; Ptak et al. 2005; Coop et al. 2008; Hinch et al. 2011; Auton et al. 2012; Stevison et al. 2016; Alleva et al. 2021; Damm et al. 2022). In these species, hotspot localization is controlled by the PRDM9 protein (PR/SET domain containing protein 9), a histone methyltransferase that specifies recombination hotspot positions (Baudat et al. 2010; Myers et al. 2010; Parvanov et al. 2010; Hoge et al. 2024; Raynaud et al. 2025). Its activity relies on four functional domains (KRAB, SSXRD,PR/SET, and a C2H2 ZnF) (Baker et al. 2017; Imai et al. 2017; Thibault-Sennett et al. 2018) among which the Zinc Finger (ZnF) domain binds to a specific DNA motif (Oliver et al. 2009; Billings et al. 2013). After binding, PRDM9 trimethylates H3K4 and H3K36 on adjacent nucleosomes through its PR/SET domain (referred to as the SET domain hereafter), creating an unique epigenetic signature that promotes DSB formation (Hayashi et al. 2005; Baudat et al. 2010; Wu et al. 2013; Eram et al. 2014; Powers et al. 2016). PRDM9 also acts in association with HELLS to access chromatin and direct the recombination machinery to its binding (Imai et al. 2020; Spruce et al. 2020). Functional PRDM9 is associated with three molecular features. First, PRDM9 needs its four domains to be intact to specify recombination hotspots (Baker et al. 2017; Imai et al. 2017; Thibault-Sennett et al. 2018) . Second, the three catalytic tyrosines (Y276, Y341, and Y357) of the SET domain are required for methyltransferase activity and are conserved in many vertebrates (Wu et al. 2013; Baker et al. 2017; Diagouraga et al. 2018; Raynaud et al. 2025). Third, in species with a functional PRDM9, the ZnF domain is fast-evolving, in particular at amino acids (AAs) directly involved in DNA binding (-1, +2, +3 and +6 positions). Multiple alleles also often coexist within populations, enabling recognition of diverse DNA motifs and driving rapid turnover of recombination landscapes (Buard et al. 2009; Capilla et al. 2014; Vara et al. 2019; Alleva et al. 2021; Raynaud et al. 2025). These properties of the ZnF domain are linked to two key evolutionary processes. First, PRDM9 binding sites undergo erosion due to biased gene conversion during DSBs repair (Myers et al. 2010; Lesecque et al. 2014; Baker, Kajita, et al. 2015). Second, the ZnF array exhibits exceptionally high diversity (Buard et al. 2009; Berg et al. 2010; Kono et al. 2014; Schwartz et al. 2014; Alleva et al. 2021; Damm et al. 2022; Raynaud et al. 2025), resulting from rapid evolution driven by a Red Queen dynamics, a co-evolutionary process in which entities constantly innovates (here the ZnF array) in reponse to the evolution of its interacting partner (here the genomic targets). This positive selection favors the emergence of new ZnF variants that recognize novel DNA binding motifs thereby countering the erosion of hotspots. Thus, the rapid evolution of the ZnF array of PRDM9 is likely related to its role in recombination (Baker et al. 2017). Beyond sequence-level diversity, *Prdm9* also displays extensive structural dynamism at the genomic level, with frequent duplication events leading to the coexistence of full-length and truncated paralogs within species. In some cases, multiple full-length copies may be present, although only one retains functional characteristics (Baker et al. 2017; Raynaud et al. 2025).

PRDM9 is part of the PRDM multigenic family, characterized by a SET domain followed by a variable number of ZnF. This multigenic family originated in metazoans and is highly dynamic (Fumasoni et al. 2007; Vervoort et al. 2015), with many expansion and rearrangements that contributed to functional diversification of the different members (Fumasoni et al. 2007; Hohenauer and Moore 2012; Di Zazzo et al. 2013; Di Tullio et al. 2022).

Despite its important role in recombination, *Prdm9* has been completely or partially lost (*i*.*e*. complete or total loss of at least one of the four domains) in many lineages (Baker et al. 2017). Partial losses are often associated with a ZnF domain that lacks signatures of rapid evolution, and recombination hotspots are then redirected toward transcription start sites or CpG islands, as shown for the swordtail fish lacking KRAB and SSXRD domains (Baker et al. 2017). Total losses of *Prdm9* are often associated with the loss of *Zcwpw1, Zcwpw2*, and, more occasionally, *Tex15* and *Fbxo47* in vertebrates (Cavassim et al. 2022). Whether this co-occurrence pattern is a feature that predates vertebrate radiation is unknown.

While PRDM9’s function has been well characterized in mammals and, to a lesser extent, in other vertebrates, its evolutionary origin and role in invertebrates remains largely unknown, even though sexual reproduction and thus meiotic recombination is a shared feature of all eukaryotes (Goodenough and Heitman 2014). Preliminary evidence, however, suggests that such a function of PRDM9 may have originate as early as it appears in Metazoans. For instance, Oliver et al. (2009) identified PRDM9 orthologs in two invertebrates, including the cnidarian *Nematostella vectensis*, and found that the ZnF domain shows signatures of positive selection at DNA-contact AAs, a functional feature of PRDM9 in vertebrates. These findings indicate that key molecular features associated with PRDM9 function could be ancestral and have been conserved in vertebrates.

A central question is thus whether PRDM9 function and its molecular features are common to all metazoans. Cnidarians are particularly relevant in this context, as they diverged early among eumetazoans and are sister to bilaterians. This makes them a valuable source of information about ancestral metazoan states.

To this end, we searched for PRDM9 in 28 species of cnidarians for which genomes and/or annotated proteomes are available. If PRDM9 plays a comparable role in cnidarians to that observed in vertebrates, then we would expect: (i) full length copies containing the four functional domains, (ii) conservation of catalytic tyrosines in the SET domain, (iii) a polymorphic ZnF domain with diversity restricted to AAs in contact with DNA.

We argued that ancestral features of PRDM9 function should be associated with the molecular features observed in vertebrates. We thus analyzed the genomes of 28 cnidarians, identifying all PRDM9 paralogs and characterizing both partial and full-length PRDM9 to reconstruct their phylogenetic relationships, assess domain architecture, conservation of catalytic residues, and ZnF diversity. These analyses provide a framework to assess whether cnidarian PRDM9 retains the molecular features linked to its canonical function and, more broadly, to clarify the gene’s evolutionary history across eumetazoans.

## Results

### PRDM9 is present in Cnidaria and shares vertebrate-like functional features

#### Genome-wide identification of *Prdm9* in 28 Cnidaria species

Among the 28 species of Cnidaria that were examined *in silico*, we identified a total number of 109 *Prdm9* homologs. Among them, 45 were full-length (FL) homologs (*i*.*e*. harboring the 4 canonical domains: KRAB, SSXRD, PR/SET and ZnF array) and 63 were truncated (T) homologs (Fig.1, Supplementary Table S5).

**Fig. 1.**
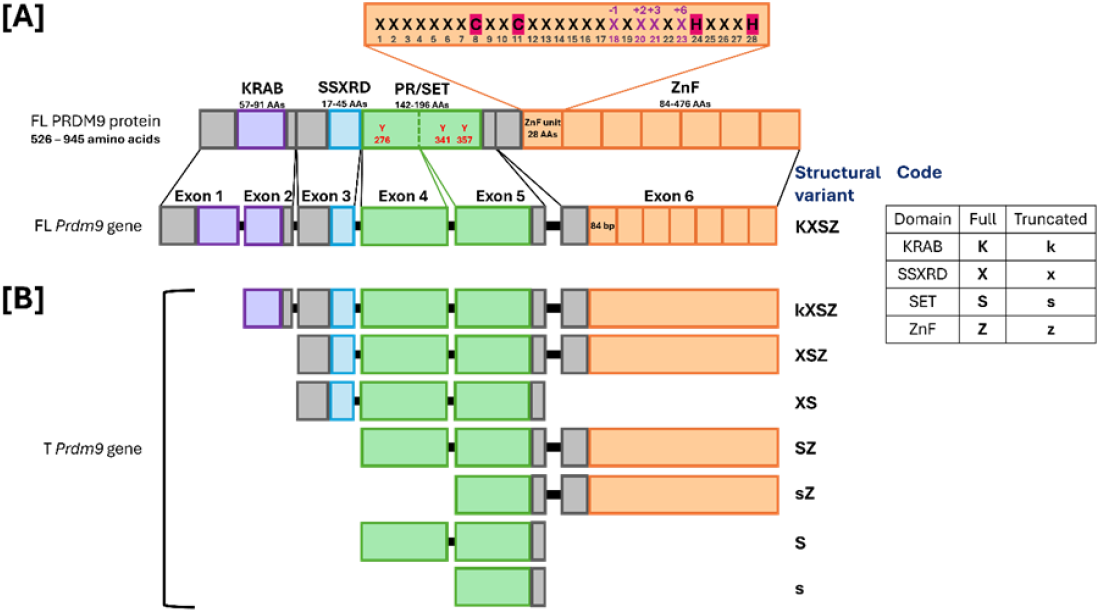
Representation of the exon structure of PRDM9 FL and T variants. Each structural variant is associated with a letter code indicating the presence of complete domains (uppercase letters) or truncated domains (lowercase letters). The KRAB (K) and PR/SET (S) domains are encoded by two exons, whereas the SSXRD (X) and ZnF (Z) domains are each encoded by a single exon. The KRAB and SET domains may be truncated due to the absence of the first exon encoding these domains. The grey boxes indicate coding sequences flanking the annotated domains and introns are represented by a black line.

FL-homologs were identified in all species (Fig. 2). However, the result for *Exaiptasia diaphana* remained questionable: a variant lacking the KRAB domain was identified in the older genome assembly, whereas a variant lacking the ZnF array was detected in the most recent one (Supplementary Table S5). As the SSXRD and SET domain sequences were identical in the two assemblies (Supplementary Figure S2) and as the genomic synteny surrounding the *Prdm9* candidates was conserved (Supplementary Table S6), we inferred the presence of a full-length *Prdm9* in *E. diaphana*. We found a unique FL-homolog for 18/28 species, including *Rhopilema esculentum, Hydractinia symbiolongicarpus, Hydra vulgaris, Dendronephthya gigantea, Exaiptasia diaphana, Actinia tenebrosa, Acropora cervicornis, Acropora hyacinthus, Acropora muricata, Acropora millepora, Montipora foliosa, Montipora capricornis, Orbicella faveolata, Pocillopora verrucosa, Pocillopora meandrina, Pocillopora damicornis, Pocillopora acuta* and *Stylophora pistillata*. Multiple FL-homologs (from two to four paralogs) were identified for ten species distributed across all cnidarian orders, with two FL-paralogs for *Nematostella vectensis, Corallium rubrum, Paramuricea clavata, Porites lobata* and *Madrepora oculata*; three FL-paralogs for *Xenia sp*., *Acropora digitifera* and *Porites evermanni*; and four FL-paralogs for *Clytia hemisphaerica* and *Desmophyllum pertusum*. All identified FL-homologs displayed the three catalytic tyrosines, except in *M. capricornis*, for which the FL-homolog contained a Y276D substitution.

**Fig. 2.**
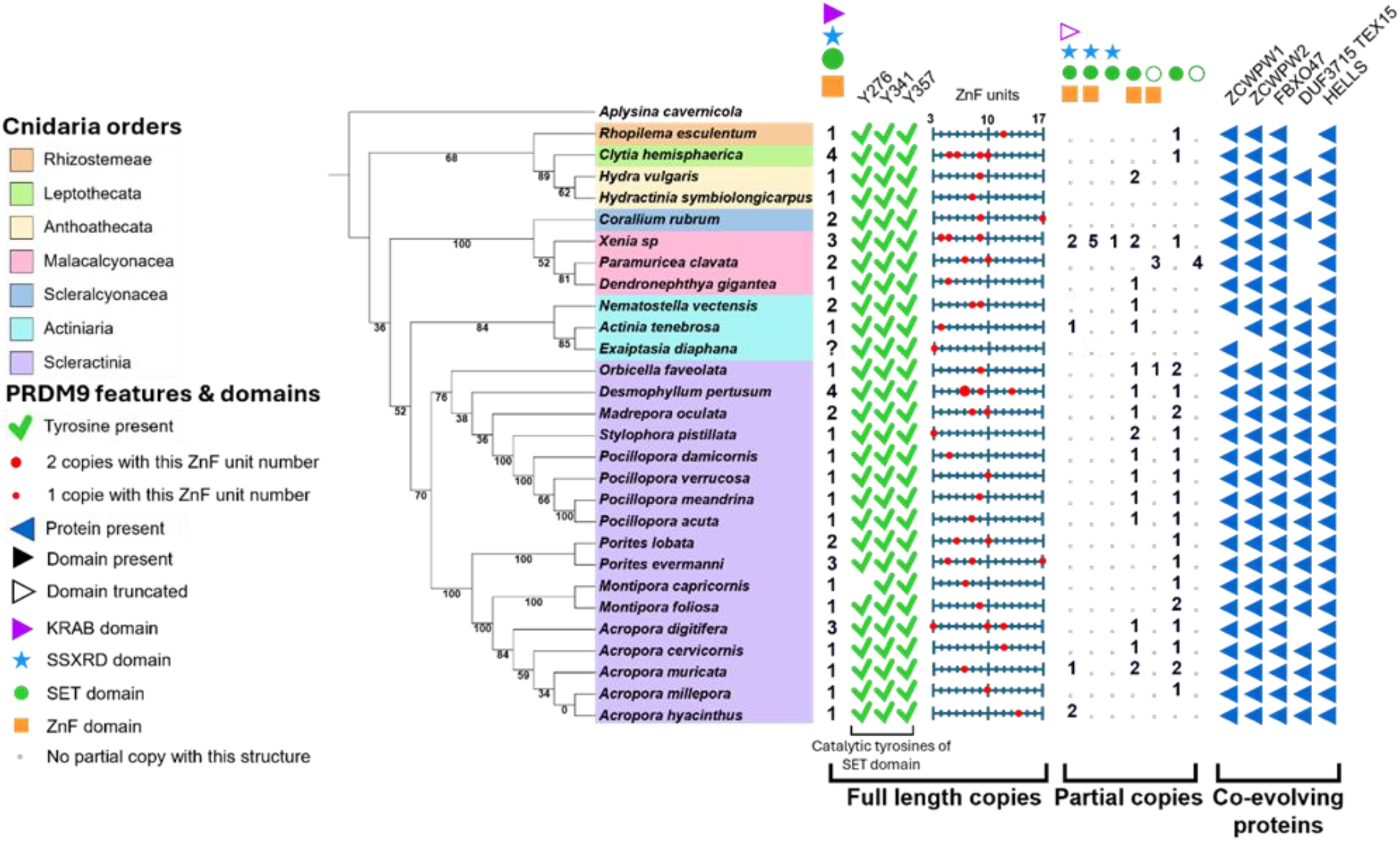
Distribution and structural diversity of full-length and truncated *Prdm9* homologs and their co-evolving proteins across 28 cnidarian species. Each PRDM9 domain is represented by a specific symbol, filled when the domain is fully present and unfilled when it is present but truncated. The KRAB domain is shown as a purple triangle, the SSXRD domain as a blue star, the SET domain as a green circle, and the ZnF domain as an orange square. For each species, the number of full-length *Prdm9* paralogs is indicated below the symbols corresponding to fully conserved domains. The “?” for Exaiptasia diaphana indicates a putative FL *Prdm9*, as a KXS variant was identified in one genome assembly, whereas an XSZ variant was found in another, at the same genomic locus. The presence of catalytic tyrosines in the SET domain is indicated by a green check mark positioned below the corresponding tyrosine. For full-length copies, the number of ZnF repeats per domain ranges from 3 to 17. The distribution of ZnF numbers is shown on a horizontal scale: a small red circle indicates that a single homolog contains n ZnFs, whereas a larger red circle indicates that two homologs contain n ZnFs; the position of the circle on the scale corresponds to the observed number of ZnFs. The architectures of partial proteins are illustrated using the corresponding domain symbols, accompanied by the number of structures identified for each species. The presence of proteins co-evolving with PRDM9, as well as HELLS, is indicated by a green check mark. Aplysina cavernicola is used as an outgroup for phylogenetic rooting.

Interestingly, in addition to these FL-homologs, we identified 63 *Prdm9* T-homologs, that have been defined as homologs containing the minimal SET domain but having lost at least one of the other domains. Their structural variations have been classified into seven categories depending on domain presence, absence and/or truncation (Fig. 1B). Structural variants all involved a 5’- and/or a 3’-end truncation and corresponded to the depletion of one or more entire exons. Consequently, domains encoded by multiple exons (*e*.*g*. two exons encoding KRAB and PR/SET) could be present and intact, present and truncated (i.e. one exon is missing), or absent (i.e. all exons are missing). Similarly, domains encoded by a single exon (*e*.*g*. SSXRD and ZnF array) could be present or absent. We encoded structural variants using four letters K, X, S, Z representing the four domains KRAB, SSXRD, PR/SET and ZnF array, respectively; uppercase was used to symbolize the full-length domain and lowercase a truncated one, no letter signifying an absence (Fig. 1B).

Not all structural variations have been observed: for example, the presence of KRAB (complete or truncated) is never associated with 3’-end losses. For structural variants consisting of two domains or fewer (*i*.*e*. SZ, sZ, S, s) we wondered whether the candidates should be classified as a *Prdm9* gene, or as a gene belonging to another PR/SET gene family, as is the case for other members of the *Prdm* family. We thus performed a phylogenetic analysis using the PR/SET protein sequence of our candidates, of identified full-length PRDM9 and of other PRDM (Vervoort et al. 2016) to identify the closest sequences. Among our 130 candidates, we excluded 79 that clustered with non-PRDM9 and assigned 51 as probable PRDM9 (Supplementary Figure S1). Among the S and s T-homologs, we often identified, using InterPro, a degenerated C2H2 ZnF domain that had lost some key residues of the characteristic X7-CXXC-X12-HXXXH motif. As both HMM and tblastn had failed to detect this domain, it was considered to be lost.

Including all possible structural variations, a total of 63 T-homologs were thus considered as *Prdm9* homologs and their catalytic tyrosines were then inspected. Most of them (44/63) conserved the three catalytic tyrosines, while others showed substitutions at one or two residues but never at the three residues (Supplementary Table S5). Structural variants kXSZ and XSZ generally preserved all three tyrosines, with a single exception in XSZ *Xenia* sp. lacking Y276.

#### Pattern of ZnF evolution in cnidarian

The number of repeated ZnF units reported from genome assemblies (Fig. 1) was highly variable between species and between variants within a species. It ranged from three (*S. pistillata, A. digitifera* and *E. diaphana*) to 17 ZnFs (*P. evermanni, P. clavata*) for FL-homologs and from 2 to 20 for T-paralogs having this domain (Supplementary Table S5). For species having multiple FL-paralogs, their ZnF arrays often have different number of ZnFs units (*e*.*g*. from 3 to 10 for the four paralogs of *C. hemisphaerica*, Fig. 2). No particular pattern of ZnF number distribution was observed when comparing species with single or multiple FL-paralogs.

As described in other metazoan species (Oliver et al. 2009; Baker et al. 2017), we noted that the four amino-acids that are contacting DNA (residues -1,+2,+3,+6 of the alpha helix) are hypervariable when comparing ZnF units in an array, whereas the other amino-acids are globally conserved across ZnFs. When comparing the proportion of diversity (PoD) carried by the four DNA-contacting amino-acids (Fig. 3), we observed that the values covered the same min-max range for FL-homologs of both vertebrates and cnidarians, from 0.33 (*Octodon degus*) to 1.0 (*Microcebus murinus, Nannospalax galili, Peromyscus maniculatus bairdii, Tupaia chinensis*) for vertebrate species (Baker et al. 2017) and from 0.29 (*Stylophora pistillata*) to 1.0 (*Acropora cervicornis, A. digitifera, A. millepora, Dendronephthya gigantea*) for cnidarians (this study), without significant difference between vertebrate and cnidarian FL-homologs (p-value = 1).We did not identify a specific pattern for the values of PoD when comparing species with single or multiple FL-homologs, as well as when comparing it between the different FL-paralogs within a species. When analyzing PoD for T-homologs, we noted a marked shift toward lower values and a narrower distribution compared to FL-homologs, in both vertebrates and cnidarians. In vertebrates, PoD ranged from 0.158 (*Clupea harengus*, SZ) to 0.524 (*Myotis lucifugus*, XSZ), whereas in cnidarians PoD ranged from 0.202 (*Hydra vulgaris*, SZ) to 0.784 (*Xenia* sp., kXSZ). Among species harboring multiple T-homologs, PoD values were variable, with no apparent intra-species pattern.

**Fig. 3.**
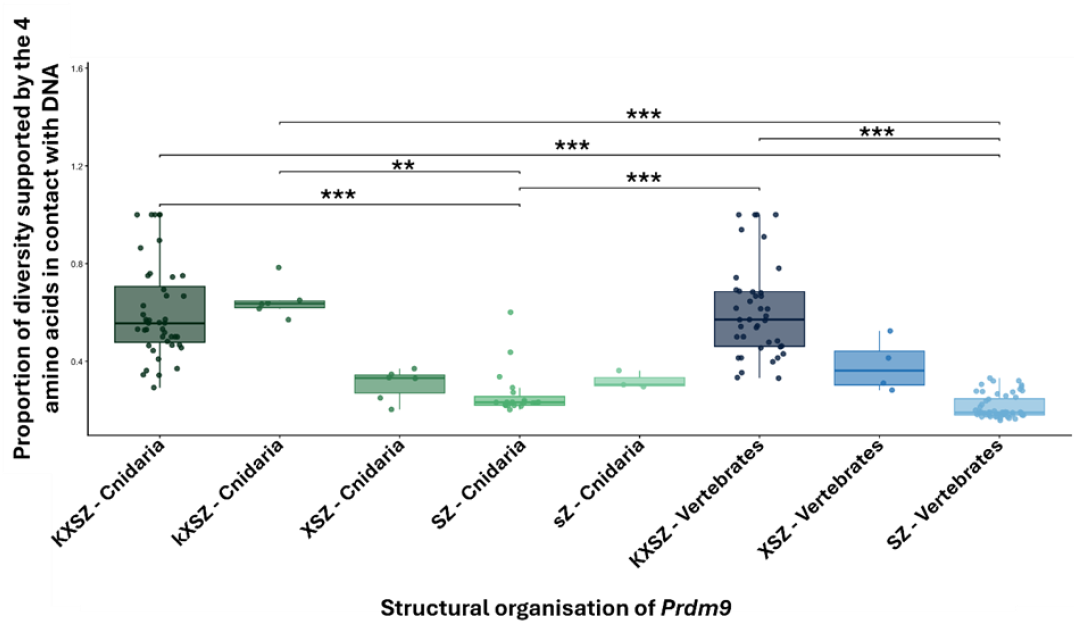
Proportion of diversity (PoD) supported by the four DNA-contacting amino acids across different PRDM9 structural architectures in cnidarians and vertebrates. Distributions are shown as boxplots for different PRDM9 architectures, distinguishing full-length proteins from partial forms characterized by the presence, absence, or partial loss of the KRAB, SSXRD, SET, and ZnF domains. Vertebrate values from Baker et al. (2017) are shown in blue, and Cnidarian values in green. Darker shades indicate full-length proteins, while lighter shades represent partial forms. Individual points correspond to single observations. Statistical comparisons between groups were performed using Dunn test with Bonferroni correction and are indicated by horizontal bars, with significance levels as follows: * p ≤ 0.05; ** p ≤ 0.01; *** p ≤ 0.001.

The distributional shift between FL- and T-homologs was primarily associated with specific structural variants. First, in both vertebrates and cnidarians, SZ T-homologs were predominantly distributed toward lower values of PoD compared to FL-homologs (cnidarians FL *vs*. SZ : p = 1.54 × 10^−5^; FL vertebrates *vs*. cnidarian SZ: p = 1.28 × 10^−5^; vertebrates FL *vs*. SZ: p = 1.45 × 10^−18^, Dunn test). Then, among T-homologs, the structural type kXSZ, identified here in cnidarians, but not yet described in vertebrates, displayed some of the highest PoD values observed among T-homologs. The distribution for kXSZ is even not different from the one of FL-homologs (p=1), but from SZ structural variants (vertebrates: p = 2.71 × 10^−6^; cnidarians: p = 5.48 × 10^−3^). Variants of type XSZ (and also sZ, for cnidarians) have PoD values that span the lowest extremes of the distribution shown for FL-homologs, but no statistical differences were identified, likely due to limited sample sizes.

Among species with relatively low PoD for FL-homologs, contrasting patterns were observed: in *N. vectensis*, its T-homolog kXSZ displayed a higher PoD (0.532) than its FL-paralog (P=0.362), whereas in *S. pistillata*, both FL- and the two SZ T-homologs consistently showed low values of PoD, ranging from 0.23 to 0.292.

### *Conservation of* Prdm9 *co-evolving genes in cnidarians*

The co-evolving *Prdm9*-associated genes *Zcwpw1, Zcwpw2, Fbxo47* and *Tex15*, together with the chromatin remodeler *Hells*, were investigated across the same 28 cnidarian genomes (Fig. 2, Supplementary Table S7). *Zcwpw1, Zcwpw2, Fbxo47* and *Hells* were identified in nearly all species, with only two exceptions: *A. tenebrosa*, which lacks *Zcwpw1*, and *E. diaphana*, which lacks *Zcwpw2*.

In contrast, *Tex15* showed a more variable pattern. The vertebrate protein typically comprises a DUF3715 domain and one or two additional conserved domains. In cnidarians, only the DUF3715 domain was detected in 21 species, which we therefore classified as truncated *Tex15* homologs. The remaining seven species (*R. esculentum, C. hemisphaerica, H. symbiolongicarpus, Xenia* sp., *D. gigantea, P. clavata* and *A. digitifera*) show a complete loss of *Tex15*.

### *History of* Prdm9 *duplications and losses in Cnidaria*

A phylogenetic analysis of sequences of all *Prdm9* homologs, based on the alignment of the SET domain (the only domain of *Prdm9* gene that was in common across all structural variants), indicates that this gene was present in full-length in the common ancestor of cnidarians (Fig. 4). The tree topology highlights several duplication and loss events that probably arose at different evolutionary timescales.

**Fig. 4.**
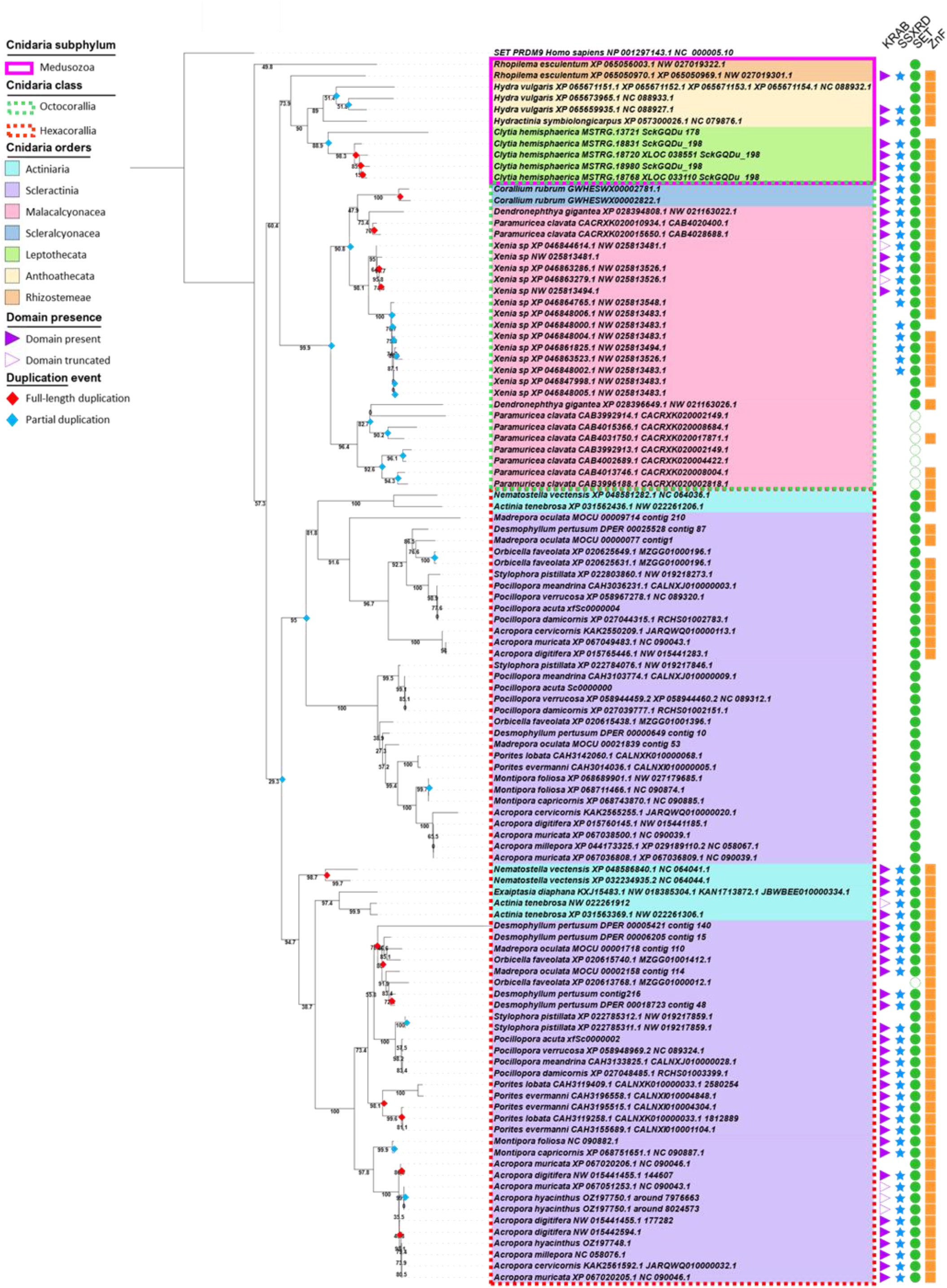
(Previous page) SET domain phylogeny for reconstruction of the history evolution of *Prdm9* in Cnidaria. Phylogenetic tree constructed with PhyML under the GTR+F+I+G4 model with 1000 bootstraps, based on the nucleotidic sequences of SET domain for FL-homologs and T-homologs from Cnidarians. Branch labels indicate the species name, the protein accession number (when available), and the corresponding nucleotide accession number. Medusozoa are highlighted in pink, Octocorallia and Hexacorallia are shown in the green and red dotted boxes, respectively, and cnidaria orders are indicated by the different colors: Rhizostemeae (orange), Anthoathecata (yellow), Lepthothecata (green), Malacalcyonaceae (pink), Sclearlcyonaceae (blue), Actiniaria (purple) and Scleractinia (cyan). Columns on the right show presence (filled) or partial loss (empty) of PRDM9 domains. Putative duplication events are indicated by a red or blue diamonds at the relevant nodes, according to whether the event gave rise to a FL or a T-homolog, respectively.

One key finding is that the multiple copies of full-length paralogs resulted mainly from recent duplication events for all taxonomic groups (see red dots located deeply in the nodes of the phylogenic tree, Fig. 4). These duplication events probably occurred after the species radiation within the Medusozoa (*C. hemisphaerica*) and the Octocorallia (*C. rubrum, Xenia sp*.and *P. clavata*) taxonomic groups, as evidenced by the clustering of these multiple FL-paralogs by species in the phylogenetic tree (Fig. 4). In Hexacorallia, many duplication events also occurred after species radiation (*e*.*g*. in *N. vectensis, D. pertusum, S. pistillata*, or *A. tenebrosa*; non-exhaustive list). However, we identified an earlier origin for some FL-paralogs. For instance, an additional FL-paralog was acquired by deep water corals (*D. pertusum, M. oculata, O. faveolata*), but was then partially lost in *O. faveolata*. Genus-specific duplications were also observed in *Porites* and *Acropora* species.

Conversely, the duplication events that gave rise to the contemporary T-homologs occurred at different timepoints in the course of cnidaria evolution (see blue dots located more or less deep at node of the phylogenic tree, Fig. 4). Our results demonstrated an early duplication in the common ancestor of Hexacorallia that gave rise to SZ paralog, that is well-conserved but lost in *Porites, Montipora* and one *Acropora* species. This SZ paralog was itself duplicated in Scleractinia only and conserved in all the species studied in this Order as a homolog with the structure S. Another duplication in the common ancestor of *P. clavata* and *D. gigantea* gave the SZ, sZ and s homologs, whose highly divergent structure could indicate a progressive loss of these copies. The earliest T-homologs, most of which arose from intra-species duplication, had a greater number of conserved domains. For example, all kXSZ and XSZ homologs emerged within a species, or within a genus, as shown by their position within terminal branches (Fig. 4).

Overall, the phylogeny of all *Prdm9* full-length and truncated paralogs highlighted an evolutionary history marked by several complete or partial duplication events that have occurred during cnidaria evolution, with several independent partial or complete losses, leading to the coexistence of complete and partial forms of the gene within almost all species studied. Our results revealed that FL homologs arose from recent events, whereas T-homologs emerged at different times, probably through long-standing exaptation for some of them.

## Discussion

Our study of *Prdm9* in Cnidaria builds upon the earlier observations by Oliver *et al*. (2009) who primarily described the rapid evolution of *Prdm9* zinc fingers across various metazoan species, including in the sea anemone *Nematostella vectensis* (Cnidaria). Subsequent literature in vertebrates has supported the idea that the fast evolution of the ZnF domain is indicative of the function of *Prdm9* in localizing recombination hotspots. Cnidaria, one of the most early diverged clades within Metazoa, are thus an ideal model to further investigate the origin of PRDM9 function. By analyzing the genomes of 28 cnidarian species, we identified at least one full-length *Prdm9* paralog in each species, all of them having the previously described molecular features indicative of PRDM9 functionality. Our results also provide the first evidence that multiple full-length paralogs, which may be functional, could coexist in the genome of at least a third of these species. Nevertheless, phylogenetic analysis revealed important birth-and-death dynamics of *Prdm9* in Cnidaria. Multiple full-length homologs originated from recent duplication events and some truncated *Prdm9* homologs, whose functions remain to be determined, probably arose from long-standing exaptations.

### Cnidarian PRDM9 share molecular hallmarks of functional vertebrate PRDM9

The 28 cnidarians analyzed here have at least one full-length *Prdm9* homolog, that is, with the four intact characteristic functional domains: KRAB, SSXRD, SET and ZnF (Grey et al. 2018; Paigen and Petkov 2018). The unique result that was not fully conclusive was related to *E. diaphana*, for which a truncated homolog missing the KRAB domain (XSZ) was identified in one assembly but a truncated homolog missing ZnF (KXS) was identified at the same locus in the most recent assembly (Supplementary Tables S1 and S5). We suggested that absence of ZnF in the most recent assembly was probably due to an assembly artefact, leading to partially reconstructed or missing genes (Florea et al. 2011; Tørresen et al. 2019). This hypothesis is supported by our own observations: in *Pocillopora acuta*, we first identified a truncated homolog (KXS form) when using one assembly (Vidal-Dupiol et al. 2020), but a full-length homolog when using a most recent one (Stephens et al. 2022). In this particular case, it is interesting to note that the ZnF domain was also missing. This could be easily explained by the minisatellite structure of ZnF domain, that probably leads to assembly errors more frequently than other domains. Independent losses of the *Prdm9* gene within specific clades have also been reported many times (Oliver et al. 2009; Baker et al. 2017; Raynaud et al. 2025). We would not have been surprised to identify a species without any full-length *Prdm9*. An alternative hypothesis could be the existence of structural variation between the two different strains of *E. diaphana* that have been used for each assembly (the F003 female strain for GCA_056151815.1 and the CC7 male strain for GCF_001417965.1), although this specific pattern has never been reported for *Prdm9* to date. This could reflect an ongoing loss of *Prdm9* in this species, a hypothesis even supported by the low diversity found at ZnF domain residues in contact with DNA (0.203, with only 3 ZnF units identified) and the absence of ZCWPW2 (see below). We thus could not completely rule out the absence of full-length homolog without additional data, for example gonad-specific long-read RNA sequencing or *de novo* assembly of other strains.

Each of these full-length homologs displayed the molecular hallmarks that have been highlighted in vertebrates and associated with the function in hotspot localization. First, the three catalytic tyrosine residues Y276, Y341 and Y357, which are conserved in all vertebrates with an full-length homolog (Baker et al. 2017), were also conserved in all full-length homologs of cnidarians, including those present in multiple copies. The sole exception is the Y276D substitution found in the unique full-length homolog of *Montipora capricornis*. These three residues should all be important for the methyltransferase activity of the protein, as shown for the mouse PRDM9 (Wu et al. 2013). However the residue Y276 is probably dispensable (Hayashi et al. 2005), and only the role of the Y357 residue has been demonstrated *in* vivo (Diagouraga et al. 2018). Also, Y357 is the most conserved residue in SET domain of PRDM proteins (Vervoort et al. 2015). Therefore, the cnidarian full-length homologs should thus be able to catalyze both H3K4me3 and H3K36me3, the unique combination of histone modifications required to direct the recombination machinery to PRDM9-binding sites. All the cnidarian full-length homologs also displayed a fast-evolving ZnF domain, as shown by the diversity concentrated at the four residues (position -1, +2, +3, +6 of the alpha helix) in contact with DNA. This ZnF diversity has been extensively described in vertebrates (Oliver et al. 2009; Buard et al. 2014; Kono et al. 2014; Baker et al. 2017; Damm et al. 2022; Hoge et al. 2024; Raynaud et al. 2025) but only sporadically in invertebrates metazoans (Oliver et al. 2009; Ponting 2011), although the cnidarian *N. vectensis* had been inspected in Oliver’s work. The rapid evolution of ZnF binding specificity is an additional hallmark of PRDM9 that indicates strong positive selection in that domain (Ponting 2011). This has been experimentally shown to result from PRDM9-directed recombination in primate and rodent species, leading to hotspot erosion (Myers et al. 2010; Lesecque et al. 2014; Baker, Kajita, et al. 2015; Davies et al. 2016; Smagulova et al. 2016). The same evolutionary force was likely driving the diversity seen in these residues in cnidarians. An extended description of PRDM9 zinc finger in natural populations of cnidarians could help us better understand the extent of this diversity and its evolutionary dynamics, particularly between pairs of paralogs.

We here also conducted the first extensive exploration of the *Zcwpw1, Zcwpw2, Tex15*, and *Fbxo47* in cnidarian genomes. In fact, *Zcwpw1* and *Zcwpw2*, and to a lesser extent *Tex15* and *Fbxo47*, co-occur (or are jointly lost) with *Prdm9* in vertebrates (Mahgoub et al. 2020; Wells et al. 2020; Cavassim et al. 2022). We reasoned that this co-evolution may predate the emergence of vertebrates, particularly in the case of *Zcwpw1* and *Zcwpw2*. Both proteins recognize the unique combination of H3K4me3 and H3K36me3 modifications catalyzed by *Prdm9* (Huang et al. 2020; Mahgoub et al. 2020; Wells et al. 2020; Cavassim et al. 2022), that is, to our knowledge, a feature restricted to *Prdm9*. Our data supported globally this model as almost all the 28 cnidaria species have both *Zcwpw1* and *Zcwpw2*, which is again a strong support of *Prdm9* functionality in this clade. However, two species have lost one of the two *Zcwpw1/2* paralogs: *E. diaphana* (lacking *Zcwpw2*) is the species for which the presence of a full-length *Prdm9* has not been firmly established, and *Actinia tenebrosa*, another sea anemone lacking the *Zcwpw1* but with a complete *Prdm9*. This pattern was observed in some species distributed across different vertebrate clades (Cavassim et al. 2022). It is therefore not excluded that the function of the missing paralog is ensured by the other paralog in some species, or for example by another CW domain–containing protein. To date, functional studies are restricted to mice and have described that ZCWPW1 is involved in repair (Huang et al. 2020; Mahgoub et al. 2020; Wells et al. 2020) and ZCWPW2 in the regulation of meiotic transcription through the recruitment of ZCWPW1 (Ruan et al. 2026) and/or the recruitment of DSB machinery at PRDM9-bound sites (Hoge et al. 2024). In the same vein, the function of both *Fbxo47* and *Tex15* has been poorly explored outside mammals and could be indirectly linked to *Prdm9* function (Cavassim et al. 2022). We found *Fbxo47* in all species, and only the DUF3715 domain of *Tex15* ortholog in 21/28 cnidarians. In fact, none of the two additional conserved domains of the vertebrate *Tex15* ortholog can be identified in cnidarians, confirming that these domains are vertebrate-specific (Schöpp et al. 2023). We also included the investigation of the presence of the gene encoding HELLS, a protein that forms a pioneer complex with PRDM9 to facilitate access to its binding sites by increasing chromatin accessibility (Imai et al. 2020; Spruce et al. 2020). In absence of HELLS, meiotic progression is impaired and PRDM9 can’t direct recombination at its binding sites (De La Fuente et al. 2006; Zeng et al. 2011; Imai et al. 2020; Spruce et al. 2020). *Hells* was not identified in Cavassim’s work as a coevolving candidate in vertebrates (Cavassim et al. 2022). Its absence in *Caenorhabditis elegans* and *Drosophila* melanogaster (Funabiki et al. 2023), two invertebrate species without *Prdm9*, had let us to make the hypothesis of a possible joint loss of *Hells* and *Prdm9* in invertebrates. Also, we found *Hells* in all species, together with *Prdm9*. Even if all these results correlate with previous work (Cavassim et al. 2022), it is worth noting that we could thus not investigate the co-evolution between *Prdm9* and *Zcwpw1, Zcwpw2, Tex15* and *Fbxo47*, but also with *Hells*, just because no lost of *Prdm9*could support this hypothesis in cnidarians. Extending this research to other clades is thus essential to understand the tight evolutionary connection between *Prdm9* and these other genes.

### Cnidarian genomes encode multiple PRDM9 paralogs likely to be functional

Our findings showed for the first time that full-length paralogs of *Prdm9* with features associated with functionality can coexist in multiple copies within the same species. To date, all PRDM9 species described in vertebrates had only one paralog that exhibits all the molecular features compatible with directing recombination hotspots (Baker et al. 2017; Cavassim et al. 2022). When multiple full-length paralogs were found, only one retained all features. This is the case in teleost fish, which typically have two full-length paralogs, only one of these having a rapidly evolving ZnF domain characterized by high diversity at DNA-contacting positions. The other paralog showed reduced variability at these sites and was therefore thought to have lost its role in hotspot localization as shown in *Oncorhynchus mykiss* (Raynaud et al. 2025). In theory, if the multiple full-length paralogs are all expressed in meiocytes, they could all contribute to the localization of recombination hotspots, thereby increasing the number of potential targets for the recombination machinery. This should function in the same way as it does for heterozygous individuals (Smagulova et al. 2016; Raynaud et al. 2025), where the proportion of recombination sites determined by each allele is variable. It was suggested that PRDM9 multimer formation directs recombination initiation at higher affinity PRDM9 binding sites in heterozygotes (Baker, Petkova, et al. 2015). Species with multiple full-length paralogs could in theory have up to eight different PRDM9 isoforms, as is the case with *Clytia hemisphaerica* or *Desmophyllum pertusum* with four full-length paralogs. If all of them are truly involved in hotspot localization, a complex molecular interplay could directly impact DSB distribution and repair, with possible homo- and heterodimer formation. It could now be worth determining how multiple full-length paralogs activity is regulated at the molecular level and what could be the consequences of their activity on recombination landscapes.

PRDM9 function has been shown to be dosage sensitive in vertebrates (Flachs et al. 2014; Baker, Petkova, et al. 2015), with negative consequences on hotspot number and meiotic progress. According to the gene dosage hypothesis (Kondrashov and Kondrashov 2006), the fixation of additional copies of dosage sensitive genes can be driven by positive selection to increase the amount of gene product. Duplicated genes also provide an ideal substrate for the fine tuning of gene dosage, for example between sexes (Gallach et al. 2011; Wyman et al. 2012). The presence of multiple copies of the *Prdm9* gene could therefore be the result of an adaptive process (Kondrashov 2012; Qian and Zhang 2014). The molecular mechanisms involved in the regulation of these duplicated genes (Veitia et al. 2008) still need to be explored in this particular case.

### Multiple truncated PRDM9 paralogs are conserved in Cnidaria

Multiple trunctated paralogs have been identified in all species except *C. rubrum, H. symbiolongicarpus* and *E. diaphana*, in varying numbers ranging from one to eleven, and with different levels of structural organization and conservation. Similar expansions of *Prdm9* genes were previously described in vertebrates (Baker et al. 2017; Raynaud et al. 2025) and more generally within the PRDM gene family (Fumasoni et al. 2007; Vervoort et al. 2015). Some of the SZ paralogs result from ancient duplications, predating the divergence between Actiniaria and Scleractinia. The fact that these ancient SZ paralogs are well conserved in most species of these clades implies that they acquired a specific function. These truncated paralogs have a SZ structure that is shared with other PRDM (Fumasoni et al. 2007; Vervoort et al. 2015) and conserved the catalytic tyrosines that are also present in the SET domain of PRDM proteins. As PRDM proteins have been described as having a role in the activation or repression of transcription (Hohenauer and Moore 2012; Di Zazzo et al. 2013; Di Tullio et al. 2022), it is possible that at least some of these PRDM9 truncated paralogs acquired similar functions.

The six identified kXSZ paralogs conserved both catalytic tyrosines and fast-evolving ZnF. As the KRAB domain is essential for PRDM9 to direct recombination hotspots (Imai et al. 2017), we conclude that fast-evolving ZnF is probably more a remnant of past activity instead of a true indication of contemporary function. This was also observed in other species, with the same conclusion (Raynaud et al. 2025). Other KRAB-ZnF proteins are under positive selection at the same AAs positions as *Prdm9* ZnF (Schmidt and Durrett 2004) and partial loss of KRAB domain is associated with an initial functional modification (Urrutia 2003). It is thus not excluded that these homologs have acquire new functions, probably recently and independently within each species, as shown by their close proximity with respective full-length paralog (Figure 4, and next paragraph).

### *Evolutionary dynamics of* Prdm9 *duplication*

The *Prdm9* duplication dynamics in cnidaria was similar to those described for ZnF C2H2 and PRDM gene families and followed a birth-and-death model (Tadepally et al. 2008; Vervoort et al. 2016). Our findings supported a single origin of full-length paralog *Prdm9*, with multiple full-length length copies that are all the result of recent duplications, mostly intra-species or intra-genus (Figure 4). This implies that full-length duplicates tend to have a relatively short lifespan, and that in the long term only one copy remains conserved. On the contrary, truncated paralogs arose at different times, with old and recent truncated paralogs. The oldest truncated paralog identified in our study was dated before the divergence between Actiniaria and Scleractinia, that is no younger than 540 Mya (Park et al. 2012), and was conserved since then in most of the species in the orders selected for our work. Many recent truncated paralogs probably emerged directly from duplication of an oldest truncated paralog, or from an incomplete duplication of a full-length paralog (*e*.*g*. for *Clytia hemisphaerica*).

High amplification of gene families was found to be a characteristic of coral genomes, accounting for a third of the predicted genes (Noel et al. 2023). We validated this observation with *Prdm9* in corals and hypothesized that this feature could extend beyond corals to encompass all cnidarian species. In fact, in several cases, multiple paralogs were located on the same scaffold, consistent with tandem duplication events potentially generated through unequal crossing-over, as previously proposed for *Prdm9* dynamics in salmonids and ruminants (Padhi et al. 2017; Raynaud et al. 2025). Such mechanisms are known to promote rapid gene gain and loss dynamics (Cooper et al. 2007) and may explain the lineage-specific expansions observed in some cnidarians, including *Xenia* sp. Overall, this evolutionary pattern seems to follow that reported for KRAB-ZnF and PRDM gene families, which are characterized by recurrent duplications, copy number variations, differential retention of paralogs across lineages and functional diversification (Fumasoni et al. 2007; Stubbs et al. 2011; Vervoort et al. 2016).

## Conclusion

This study is the first to provide an in-depth characterization of the presence and evolution of *Prdm9* outside vertebrates, using early-diverging cnidaria as models. Our results are consistent with an ancestral origin of *Prdm9* in eumetazoans. *Prdm9* is widely conserved across several cnidarian lineages, but we can’t exclude one or more losses in cnidarian species not included in our study. We also emphasized the significant number of truncated paralogs, some of which being conserved for millions of years of evolution and providing evidence of genetic exaptation. The function of these paralogs remains to be studied. Mostly, our work is also the first to identify species with multiple full-length paralogs that have all molecular hallmarks indicative of a function in localizing recombination hotspots. This raises the question of their functional redundancy, as well as the evolutionary significance and advantage of such duplications. Regardless of how dispensable they may be, the long-term evolution seems to select back to one *Prdm9* copy. In fact, the multiple full-length paralogs are all recent, which is not the case for truncated versions. Altogether, this work demonstrates the importance of considering the full diversity of metazoans when studying the evolutionary origins of such key gene, which is involved in both meiosis and fertility.

## Materials and Methods

### In silico *identification of* Prdm9 *in cnidarians*

#### Species Selection

A total of twenty-eight Cnidaria species representing seven taxonomic orders (Rhizostomeae, Leptothecata, Anthoathecata, Malacalcyonacea, Scleralcyonacea, Actiniaria, Scleractinia) were selected for the present study. Twenty-five were chosen based on the availability of an annotated proteome obtained from assemblies at the time of the search (between October 2023 and October 2024). These proteomes were retrieved from NCBI or other databases (see Supplementary Table S1 for details). To extend the evolutionary representation within Hexacorallia and Octocorallia, we also included the following three species: *Acropora hyacinthus, Pocillopora acuta*, and *Corallium rubrum* for which genome sequences but not annotations were available.

#### Identification of *Prdm9* homologs HMM-based approach

To investigate the presence of PRDM9 in cnidarian proteomes, we searched for all candidate proteins that had one or more of the four canonical PRDM9 domains (KRAB, SSXRD, SET and ZnF array). Domain-specific profile hidden Markov models (HMM) were first generated using a multiple sequence alignment of annotated PRDM9 protein sequences from 97 species representing different clades of metazoans, ranging from porifera to primates (supplementary table S2). The hmmbuild program (version 3.1b1) of the HMMER tool suite (Finn et al. 2011) was used with the default settings on each of the four domain-restricted part of alignment. Such generated profiles capture both conserved positions and variations specific to each domain. In each of the 25 proteomes, PRDM9 domains were identified individually by homology as those matching with profiles using hmmsearch program (version 3.1b1) with the following parameters: -E = 100, --domE = 100, --F1 = 0.05, --F2 = 0.01, --F3 = 0.0001, --nobias, --nonull2. Significant matches were discriminated using bit scores (>20) and E-values (<10^−3^). Co-occurrence of domains in the same protein was inspected and the corresponding protein ID collected using the multijoin tool (Galaxy version 1.1.1) (Grüning et al. 2018). PRDM9 candidates were then determined as any protein containing one to four domains (but must always include SET), in order to retain both full-length (FL) and truncated (T) homologs (Supplementary Fig. S1).

#### Tblastn approach

As a complement of the HMM-based approach, the genome-based method tblastn (Camacho et al. 2009; Cock et al. 2015) was conducted to compare protein query sequences to translated genomic sequences (Altschul et al. 1997). Candidate genes that may be missed in the HMM approach due to incorrect exon annotation could thus be identified, as well as candidate genes in the three species with no available proteome. For each of the 28 species, we searched for all candidate genes containing one or more of the four canonical PRDM9 domains (KRAB, SSXRD, SET and ZnF array). To do that, amino-acid sequences of each PRDM9 domain were used individually as queries in the genome of the target species. The query sequences were selected based on the results of our HMM-based analysis (see above), either from the target species itself or from a phylogenetically close species but always from a full-length PRDM9 homolog (see supplementary table S1 for details). Matching genomic sequences were identified using NCBI BLAST+ tblastn (version 2.10.1+galaxy0) with the following settings: -max_target_seqs = 1000, -max_hsps = 10 and -word_size = 3 (Altschul et al. 1997; Camacho et al. 2009; Cock et al. 2015). All loci matching co-occurrence of domains were collected using the multijoin tool (Galaxy version 1.1.1) (Grüning et al. 2018). *Prdm9* candidates (FL and T) were identified as locus having one to four (but must always include SET) domains matching the same contig, with domain architecture preserved and distance between domain-encoding exons not exceeding 5kb.

#### Comparison and selection of *Prdm9* candidates

When both methods have been conducted (*i*.*e*. for 25/28 species), sequences matching the same locus were visually inspected and aligned in order to compare the number of domains as well as sequence length. When sequence length did not match, we retained the longest candidate after checking presence of start codon and absence of stop codon within the nucleotide sequence. Any candidate sequence found by tblastn at one locus but not found by HMMER (we never experienced the reverse situation) was checked for start and stop codons and added to the pool of *Prdm9* candidates. Candidates containing only SET or SET with ZnF were discriminated as either *Prdm9* homolog or other PRDM-related homolog by phylogenetic analysis based on the SET protein sequence (see details about methods below, part “Phylogenetic analysis”).

We then classified all retrieved *Prdm9* homologs according to their structural variations: full-length (FL) or truncated (T), and also refined our description based on domain composition of each homolog. A coding system based on structural variations was defined as follows: KRAB, SSXRD, SET and ZnF domains are represented by the letters K, X, S and Z, respectively. Uppercase letters indicate full-length domains, whereas lowercase letters denote truncated domains (see Fig. 1 for details). We noted that the ZnF domain could often be identified in S or s homologs using InterPro, even though neither the HMM nor the tblastn methods found it. Upon visual inspection, we concluded that the C2H2 ZnF motif X7-CXXC-X12-HXXXH had degenerated and considered the domain to be absent in these cases.

Overall, the two approaches were complementary: tblastn enabled the detection of sequences missed by HMM, often due to annotation errors into predicted proteomes (*e*.*g*. exon missing), whereas HMM provided higher sensitivity and more precise domain boundaries.

We encountered discrepancies when using one or another assembly for *Exaiptasia diaphana*: either KRAB or ZnF was missing in one but not the other assembly (Supplementary Tables S1 and S5). We thus compared the SET and SSXRD domain sequences identified in the two assemblies by performing a protein sequence alignment using MUSCLE v3.8.31 in SeaView v5.1 (Gouy et al. 2010). In addition, the local synteny was checked between assemblies by comparing similarity of the genes located upstream and downstream of the *Prdm9* locus (supplementary Fig.S2). We concluded that the locus was the same and that the gene was probably full-length, even if we could not exclude structural variation between the two strains used to produce the two different assemblies.

#### Conservation of catalytic tyrosines

In humans, the three catalytic tyrosine residues Y276, Y341, and Y357 of the PR/SET domain are essential for the methyltransferase activity of PRDM9 (Wu et al. 2013). We examined their conservation in each identified FL- and T-paralog of PRDM9 having a PR/SET domain (109 sequences) by performing amino acid alignments with the human PRDM9 sequence (NP_001297143.1) using the MUSCLE algorithm implemented in the Seaview software (Gouy et al. 2010). The conservation of the three catalytic tyrosines was visually inspected.

#### Evolution of PRDM9 ZnF domain

For each FL- or T-homolog with a ZnF domain, we reported the number of ZnF units in the domain as the number of entire ZnFs (i.e. with 28 amino-acids) matching the C2H2 ZnF motif X7-CXXC-X12-HXXXH. The amino-acid diversity of ZnF array was then inspected with a specific focus on the DNA-binding residues (position 18-20-21-23, also referred as -1, +2, +3, +6 residues relative to the alpha-helix, see Fig. 1A). For each homolog, the proportion of diversity carried by these four residues was calculated across ZnFs in the array, as the ratio between the sum of diversity at these positions and the sum of diversity across all 28 amino-acid position, as previously done by Baker et al. (2017) and Raynaud et al. (2024). This measure named PoD thus reflects the proportion of total variability carried by the residues responsible for DNA recognition, that have been described to evolve rapidly in the array of metazoans (Oliver et al. 2009).

The values of PoD obtained for different structural variants of PRDM9 in Cnidaria were compared to those previously reported in vertebrates. Statistical analyses were performed using R in RStudio software version 4.4.1 (v2024.04.2), with data handling and visualization conducted using the packages dplyr (v1.1.4, Wickham et al. 2026), ggplot2 (v4.0.1, Wickham 2016), and ggpubr (v0.6.2, Kassambara 2026), and statistical tests performed using the base stats package (v4.4.1). The normality of data distributions was assessed using the Shapiro–Wilk test. As the data did not follow a normal distribution, non-parametric statistical tests were applied. Differences in values of PoD for FL- *vs*. T-homologs and for cnidarians *vs*. vertebrates were evaluated using the Wilcoxon test. To refine this analysis, comparisons of PoD were also performed between each structural variations of T-homologs (*i*.*e*. kXSZ, XSZ, SZ, sZ). A Dunn test was then performed on these subdivisions to identify which structure differed significantly from FL-homolog in both cnidarians and vertebrates, with p-values adjusted for multiple comparisons using a Bonferroni correction. Levels of statistical significance were defined as * (p ≤0.05), ** (p ≤ 0.01), and *** (p ≤0.001).

### *Detection of genes co-evolving with* Prdm9 *and its* Hells *partner*

The four genes *Zcwpw1, Zcwpw2, Tex15 and Fbxo47*, that have been shown to co-evolve with *Prdm9* in vertebrates (Cavassim et al. 2022), and the gene *Hells*, whose protein is involved in direct interaction with *Prdm9* (Imai et al. 2020; Spruce et al. 2020), have been searched into the same 25 predicted proteomes and 28 genomes of Cnidarians. Both HMM-based and tblastn-based approaches have been conducted.

HMM profiles were generated following the procedure described above. For both *Zcwpw1* and *Zcwpw2* paralogs, one profile was built for the ZnF-CW domain and another for the PWWP domain. The reference sequences used to produce the multiple alignment were from *Zcwpw1* and *Zcwpw2* identified in different metazoan species, from porifera to human. The hmmbuild program then produced the profiles using default parameters. *Tex15* is composed of three domains: TEX15_1, TEX15_2, and DUF3715. Domain-specific profiles were constructed for TEX15_1 and TEX15_2 from the sequences collected in 14 metazoan species. As DUF3715 domain is present both in TEX15 protein and in TASOR protein, two distinct DUF3715 profiles were prepared: one from TEX15 protein sequences (DUF3715_TEX15, from sequences of 72 metazoan species) and the other from TASOR protein sequences (DUF3715_TASOR, from sequences of 77 metazoan species), to refine the origin of any DUF3715 domain that could be found alone (which is the case in cnidarians). Four profiles have thus been used to search for *Tex15*. For *Fbxo47* and *Hells*, a profile was built for each using as the reference sequences of full-length proteins from 41 and 9 metazoan species, respectively. All details regarding species and reference sequences used for each profile are available in Supplementary Table S2. Each profile was then used to scan the 25 proteomes. The matching profiles were reported for bit scores >20 and E-values <10^−3^ and the presence/truncation/absence was compiled for each candidate.

To complement this approach, we searched the five genes in the translated genome of the 28 species. The full-length protein sequence of *P. lobata*, identified using HMM profiles (see above-mentioned method) was used as query for *Zcwpw1, Zcwpw2, Fbxo47* and *Hells*, as an intermediate species in the cnidarian phylogeny. For *Tex15*, three domain-specific queries were employed: the DUF3715 domain identified in *P. lobata* and TEX15_1 and TEX15_2 from *Homo sapiens* (XP_011542890.1), as these latter domains have not been found in cnidarians.

### Phylogenetic analysis

#### Phylogenetic tree reconstruction of cnidaria species

A phylogenetic tree was reconstructed based on *Cox1* nucleotide sequences (Fig. 2). *Cox1* sequences were retrieved from mitochondrial genome annotations when available, or identified using tblastn when not annotated (Supplementary Table S3). The *Cox1* sequence of *Aplysina cavernicola* (Porifera) was used as an outgroup. Multiple sequence alignment of all *Cox1* sequences was performed using the MUSCLE version 3.8.31 implemented in SeaView version 5.1 and subsequently cleaned with Gblocks, to remove contiguous non-conserved positions (Gouy et al. 2010). Phylogenetic relationships between species were reconstructed using a distance-based Neighbor-Joining method based on pairwise non-synonymous substitution rates (Ka). The analysis was conducted on 1085 conserved sites, and branch support was assessed using 1000 bootstrap replicates.

#### Discriminating between *Prdm9* and non-*Prdm9* SET-domain-containing truncated homologs

Among the identified T-homologs candidates, we questioned whether the structural variations SZ, sZ, S and s (Fig.1B) originated from an ancestral *Prdm9* homolog or from another gene of the Prdm family, which are also characterized by the presence of both a SET and a ZnF domain. A phylogenetic analysis was conducted (Supplementary Fig. S1) using (i) the SET protein sequences from all PRDM family members identified in Vervoort et al. 2016 (Supplementary Table S4), including SET from PRDM9 of seven species (Supplementary Table S4), (ii) eight SET sequences from FL-homolog of PRDM9 identified in the present study and selected in different Classes of cnidarians (Supplementary Table S4) and (iii) the SET sequence from Human SETD6 as an outgroup (accession number : Q8TBK2; Supplementary Table S4). Multiple sequence alignment of all SET sequences was carried out with MUSCLE version 3.8.31 in SeaView version 5.1 (Gouy et al. 2010). Maximum likelihood analyses were conducted in IQ-TREE (Nguyen et al. 2015). ModelFinder selected JTTDCMut+G4 as the best-fit model under the BIC criterion (Kalyaanamoorthy et al. 2017). Branch support was estimated using 1000 bootstrap replicates (Hoang et al. 2018). The candidates T-homologs were classified to be PRDM9 if they clustered within the same clade as known PRDM9 SET sequences in the phylogeny.

#### Evolutionary history of *Prdm9* paralogs in Cnidarians

To reconstruct the evolutionary history of the multiple *Prdm9* paralogs in cnidarians, we built a phylogeny based on the nucleotide sequence of the SET domain, as this domain is the only one shared by all structural variants. Multiple alignment were obtained for the sequences of all FL- and T-homolog using MUSCLE implemented in SeaView (Gouy et al. 2010). Maximum likelihood analyses were conducted in IQ-TREE (Nguyen et al. 2015). ModelFinder selected GTR+F+I+G4 as the best-fitting model according to the Bayesian Information Criterion (Kalyaanamoorthy et al. 2017). Branch support was assessed using 1,000 bootstrap replicates (Hoang et al. 2018). The SET domain of human PRDM9 (NP_001297143.1) was used as the outgroup.

## Supporting information

SuppFig1&2

## End Matter

### Author Contributions and Notes

J.A.J.C. designed the research project. J.A.J.C. acquired fundings. L.D. provided materials and support for the methodology. A.R. collected and analysed the data. A.R. prepared figures. E.T. and J.A.J.C. supervised the work. L.D. and C.G. validated the results. A.R., E.T. and J.A.J.C. wrote the original draft ; all authors reviewed, edited and validated the manuscript.

## Acknowledgments

We thank Catriona Munro, Sophie Arnaud-Haond and Adrien Tran Lu Y for providing early access to *C. hemisphaerica, D. pertusum and M. oculata* genomes. We thank Cristian Chaparro for providing support with the local galaxy server and for helpful advises on software usage. We thank Pierre-Alexandre Gagnaire, Marie Mirouze and Catriona Munro for fruitful discussion. We thank the Coral group from IHPE, particularly Jérémie Vidal-Dupiol, Olivier Rey and Pauline Buso for weekly critical advises and comments on this work and the HotRec team for helpful feedback since the beginning of this project.

J.A.J.C group is supported by ANR [ANR-23-CE12-0011]. A.R is funded by PhD fellowship from UPVD. This study was also supported by the École Universitaire de Recherche TULIP-GS [ANR-18-EURE-0019]. IHPE lab is supported by the « Laboratoire d’Excellence (LabEx) » TULIP [ANR-10-LABX-41].

