## Supplementary material for "The eumetazoan origin of PRDM9 function revealed by cnidarian genome analysis": SuppFig1&2

### Slide 1
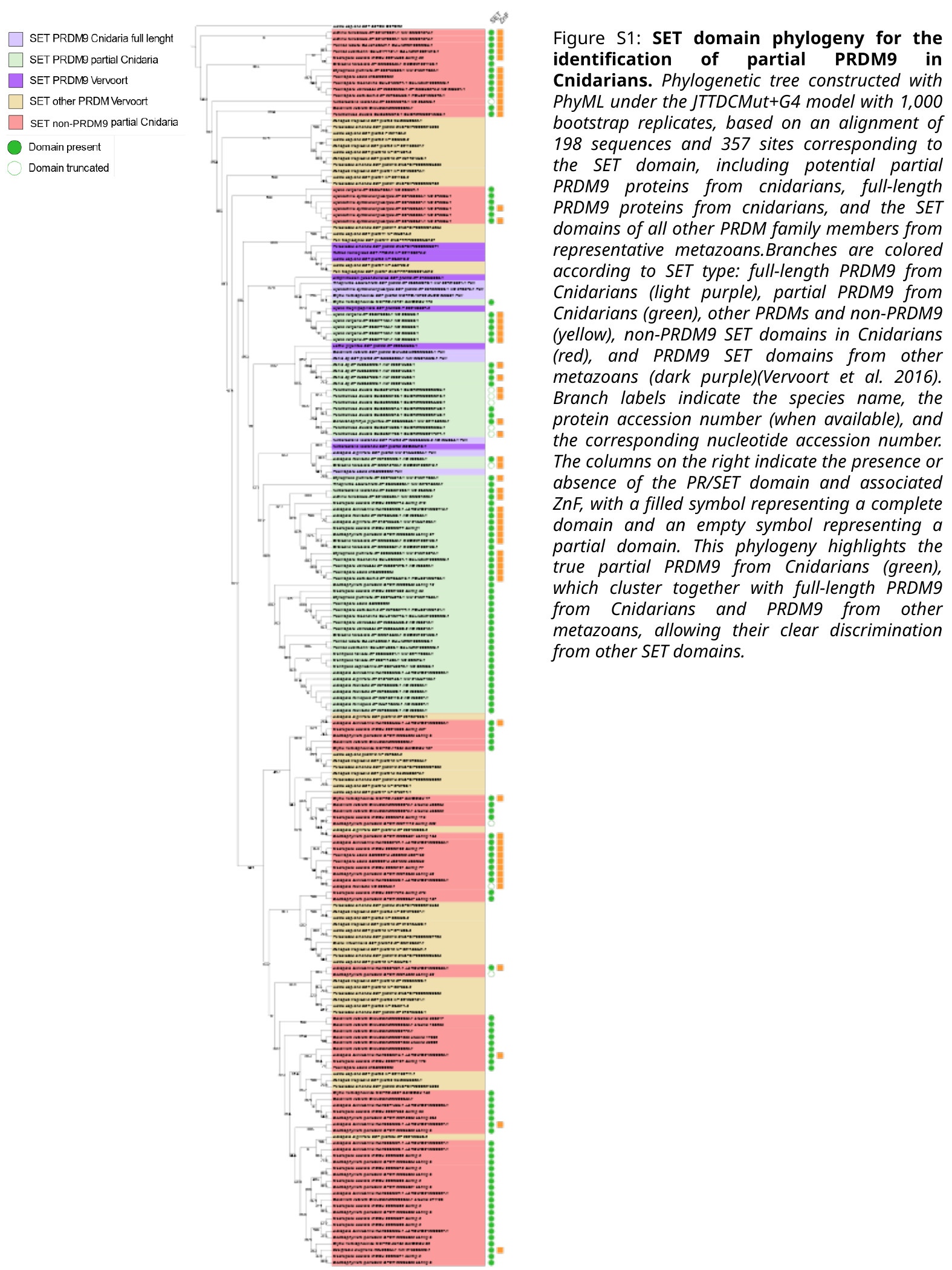

Figure S1: SET domain phylogeny for the identification of partial PRDM9 in Cnidarians. Phylogenetic tree constructed with PhyML under the JTTDCMut+G4 model with 1,000 bootstrap replicates, based on an alignment of 198 sequences and 357 sites corresponding to the SET domain, including potential partial PRDM9 proteins from cnidarians, full-length PRDM9 proteins from cnidarians, and the SET domains of all other PRDM family members from representative metazoans.Branches are colored according to SET type: full-length PRDM9 from Cnidarians (light purple), partial PRDM9 from Cnidarians (green), other PRDMs and non-PRDM9 (yellow), non-PRDM9 SET domains in Cnidarians (red), and PRDM9 SET domains from other metazoans (dark purple)(Vervoort et al. 2016). Branch labels indicate the species name, the protein accession number (when available), and the corresponding nucleotide accession number. The columns on the right indicate the presence or absence of the PR/SET domain and associated ZnF, with a filled symbol representing a complete domain and an empty symbol representing a partial domain. This phylogeny highlights the true partial PRDM9 from Cnidarians (green), which cluster together with full-length PRDM9 from Cnidarians and PRDM9 from other metazoans, allowing their clear discrimination from other SET domains.
SET non-PRDM9

### Slide 2
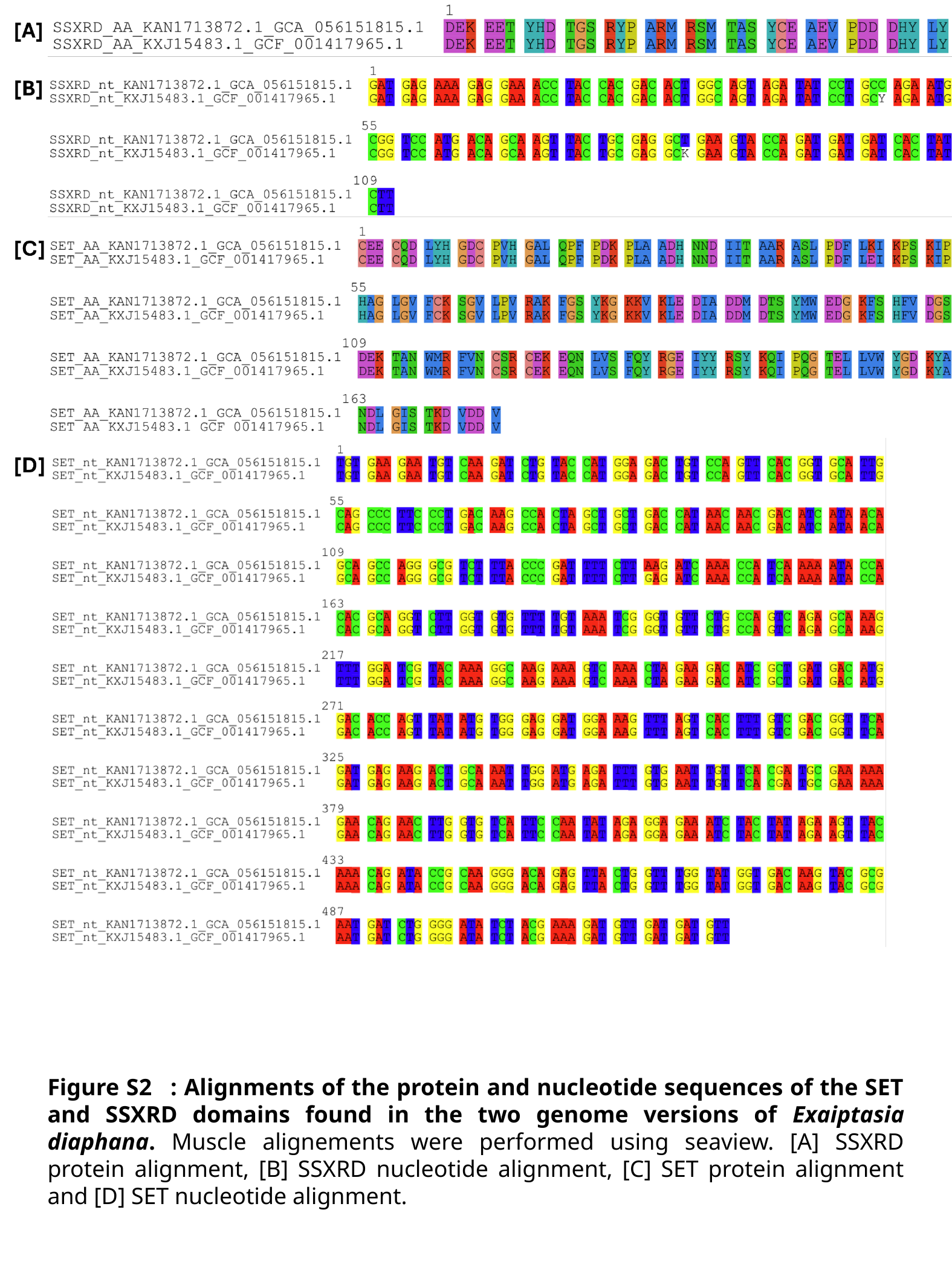

Figure S2   : Alignments of the protein and nucleotide sequences of the SET and SSXRD domains found in the two genome versions of Exaiptasia diaphana. Muscle alignements were performed using seaview. [A] SSXRD protein alignment, [B] SSXRD nucleotide alignment, [C] SET protein alignment and [D] SET nucleotide alignment.
